# PETISCO is a novel protein complex required for 21U RNA biogenesis and embryonic viability

**DOI:** 10.1101/463711

**Authors:** Ricardo J. Cordeiro Rodrigues, António Miguel de Jesus Domingues, Svenja Hellmann, Sabrina Dietz, Bruno F. M. de Albuquerque, Christian Renz, Helle D. Ulrich, Falk Butter, René F. Ketting

## Abstract

Piwi proteins are important for germ cell development in almost all animals studied thus far. These proteins are guided to specific targets, such as transposable elements, by small guide RNAs, often referred to as piRNAs, or 21U RNAs in *C. elegans*. In this organism, even though genetic screens have uncovered a number of potential 21U RNA biogenesis factors, little is known about how these factors interact or what they do. Based on the previously identified 21U biogenesis factor PID-1, we here define a novel protein complex, PETISCO, that is required for 21U RNA biogenesis. PETISCO contains both potential 5’-cap and 5’-phosphate RNA binding domains, suggesting involvement in 5’ end processing. We define the interaction architecture of PETISCO and reveal a second function for PETISCO in embryonic development. This essential function of PETISCO is not mediated by PID-1, but by TOST-1. Vice versa, TOST-1 is not involved in 21U RNA biogenesis. Both PID-1 and TOST-1 are small, intrinsically disordered proteins that interact directly with the PETISCO protein ERH-2 (enhancer of rudimentary homolog 2) using a conserved sequence motif. Finally, our data suggest an important role for TOST-1:PETISCO in SL1 homeostasis in the early embryo. Our work describes the first molecular platform for 21U RNA production in *C. elegans*, and strengthens the view that 21U RNA biogenesis is built upon a much more widely used, snRNA-related pathway.

## Introduction

Germ cells in many organisms depend critically on the integrity of a small RNA-driven pathway known as the piRNA pathway ^1 2^. This pathway is characterized by members of the Piwi protein family that, bound to their small RNA cofactors (piRNAs), act in gene-regulatory and transposon silencing pathways ^3,4^. In the absence of these pathways, germ cells do not form properly, ultimately leading to sterility. Even though this type of pathway is found in the germ cells of most, if not all animals, the mechanistic details of the Piwi pathways are remarkably different. In flies, for example, a piRNA amplification loop, driven by two Piwi paralogs, is coupled to the activity of a third, nuclear Piwi protein that drives transcriptional silencing ^5 4^. In other animals, such as the silk moth, the nuclear branch seems to be absent, while the Piwi amplification loop is present ^6^, and in mice a linear Piwi pathway drives nuclear accumulation of a Piwi protein ^4^. While transposons represent major targets for these pathways, it is also clear that non-transposon targets are functionally relevant for these pathways ^7^.

In *C. elegans*, the main Piwi protein is named PRG-1 (piwi related gene 1) and the small RNA co-factors bound by PRG-1 are known as 21U RNAs ^8-10^. This name stems from the fact that these small RNAs are 21 nucleotides long, and have a strong bias for a 5’ uracil. Target RNA recognition by PRG-1:21U complexes does not depend on full-length base-pairing between the 21U RNA and the target RNA ^11,12^, and results in the recruitment of RNA-dependent RNA polymerase activity, that drives the synthesis of so called 22G RNAs ^8,9^. These 22G RNAs, named after their predominant 22 nucleotide length and 5’ G bias, are bound by argonaute proteins that are specific for nematodes (also referred to as WAGO-proteins), that ultimately drive the silencing of the 21U target.

In absence of PRG-1, the germline deteriorates over generations, eventually leading to sterility ^13^. The defects that accumulate are most likely not genetic in nature, since the phenotype can be reverted by, for instance, starvation ^13^. These observations have led to the suggestion that in *prg-1* mutants a form of stress accumulates and ultimately leads to germ cell dysfunction ^14^. More acute fertility defects can be observed in *prg-1* mutants when the 22G RNA biogenesis machinery is reactivated in zygotes, after being defective in both of the parents. In this case, maternally provided PRG-1:21U complexes are required to prevent immediate sterility, by preventing accumulation of 22G RNA populations that inappropriately silence genes that should be expressed ^15,16^. This demonstrates that PRG-1, and its bound 21U RNA, has a critical function in maintaining a properly tuned 22G RNA population in the germ cells. Nevertheless, the impact of PRG-1 on transposon silencing is rather modest, as in *prg-1* mutants only activation of the Tc3 transposon has thus far been demonstrated ^8^.

Interestingly, the silencing of a 21U target, at least a transgenic one, can become independent of 21U RNAs themselves ^5,17,18^. In this state, which has been named RNAe (for <u>RNA</u>i induced <u>e</u>pigenetic silencing), the silencing has been completely taken over by a self-sustaining 22G RNA response. This includes a nuclear component that changes the histone methylation status of the targeted transgene, driven by the nuclear Argonaute protein HRDE-1 ^5, 17-19^. Possibly, such an RNAe state may explain why transposons are not more broadly upregulated in *prg-1* mutants, because in *prg-1;hrde-1* double mutants the Tc1 transposon is strongly activated ^15^. How exactly RNAe is established is not clear, but once it is, it can be extremely stable, for over tens or more of generations ^5,17,18^.

Given the important function of 21U RNAs driving a potentially very powerful silencing response, the biogenesis of 21U RNAs is a critical process for the germ cells of *C. elegans*. Nevertheless, very little is known about the mechanistic steps of this process. The large majority of genes encoding 21U RNAs are found in two main clusters on chromosome IV, and are characterized by a very specific sequence motif in their promoter that defines the 5’ end of the mature 21U RNA ^20^. Transcription of these genes requires a protein named PRDE-1 and the transcription factor SNPC-4 ^21,22^, the latter of which is also known to be involved in transcription of other short structural RNAs, such as snRNAs and splice-leader RNAs ^21^. An evolutionary analysis of 21U RNA loci in diverse nematodes has revealed that 21U loci may have evolved from snRNA loci ^23^. These loci include both the strongly conserved U1, U2 loci, as well as loci producing so-called splice leader RNAs (SL1 and SL2), that are trans-spliced to the 5’ ends of a large fraction of all mRNAs in *C. elegans* ^24^. These observations raise the possibility that also other aspects of the 21U RNA pathway may have mechanistic links to snRNA biogenesis.

The 21U RNA precursor transcripts are short, around 27 nucleotides, and capped ^25^. Even though a biochemical reconstitution of the processing has not been achieved thus far, the available data suggest the following order of steps in the maturation of the 21U precursor transcripts into mature 21U RNAs ^20,22,25-27^. First, the precursors are most likely processed at the 5’ end, resulting in de-capping and removal of two nucleotides. The enzymes involved have not yet been identified, and whether this reaction is mediated by endo- or exo-nucleolytic activities is not clear. This step is likely followed by loading of the 5’processed precursor into PRG-1 and trimming of the 3’ end by the 3’-5’ exonuclease PARN-1 ^28^. Finally, the 3’ end is 2’-O-methylated by HENN-1 ^29-31^. Not much is known about other proteins acting at these 21U maturation steps, even though a number of genes have been implicated in this process ^21,22,26,27 32^.

Here, we follow up on our previous identification of PID-1 (<u>p</u>iRNA induced silencing <u>d</u>efective 1) as a protein essential for 21U RNA production ^26^. Mutants lacking PID-1 produce very low amounts of mature 21U RNAs, and the 21U-related molecules that remain tend to be precursor transcripts, suggesting PID-1 acts at some step in 21U precursor processing. Other factors potentially acting at this step of 21U biogenesis, TOFU-1, 2, 6 and 7, were identified in a genome-wide RNAi screen ^27^. How these factors interconnect, however, remained unclear. We find, using immuno-precipitation (IP) and label free quantitative mass spectrometry (IP-MS), that PID-1 interacts with two proteins that were identified in a genome-wide RNAi screen for 21U RNA biogenesis factors: TOFU-6 and the unnamed protein Y23H5A.3, henceforth referred to as PID-3. In addition, we identify two strongly conserved proteins interacting with PID-1: IFE-3, a *C. elegans* eIF4E homolog, and ERH-2, one of the *C. elegans* homologs of ‘enhancer of rudimentary’. Enhancer-of-rudimentary homologs are evolutionary very well conserved proteins, with homologs being present from plants to man. Its mechanistic role is not very well established, but in *Schizosaccharomyces pombe* ERH1 drives the decay of meiotic transcripts, and interacts with the nuclear exosome and the nuclear CCR4-NOT complex ^33^. Since we always find PID-3, ERH-2, TOFU-6 and IFE-3 together in a complex we named it PETISCO, for <u>P</u>ID-3, <u>E</u>RH-2, <u>T</u>OFU-6, IFE-3 <u>s</u>mall RNA <u>Co</u>mplex. All PETISCO proteins are required for 21U biogenesis. Additionally, PETISCO mutants display a maternal effect lethal (Mel) phenotype, whereas *pid-1* and *prg-1* mutants are viable. We find that this is caused by the fact that besides binding to PID-1, PETISCO can bind a protein with similarities to PID-1. We named this protein TOST-1, for <u>T</u>wenty-<u>O</u>ne U pathway antagoni<u>st</u>. Mutants for *tost-1* produce 21U RNAs, display enhanced 21U-driven silencing and have a Mel phenotype. PID-1 and TOST-1 both interact with ERH-2 using a conserved motif, strongly suggesting a mutually exclusive mode of binding to PETISCO, where PID-1 implicates it in 21U RNA biogenesis and TOST-1 in another pathway that is essential for embryonic viability. Our data suggest that this pathway may be involved in providing the embryo with resources required for trans-splicing, especially with the SL1 splice leader RNA.

With these findings, we pave the way for a better understanding of how 21U precursor transcripts are processed. The identification of PETISCO, plus its implication in at least two distinct processes provides the new insight that 21U RNA processing is closely related to a more widely conserved process: that of snRNP biogenesis. Combined with the fact that 21U RNA transcription also bears strong signatures of an snRNA-history, an image arises in which the 21U RNA pathway may have evolved out of an already existing small non-coding RNA network that became linked to an Argonaute-driven gene-silencing program.

## Results

### Identification of PID-1 interactors

To identify proteins interacting with the 21U biogenesis factor PID-1 ^26^ we performed IPs with PID-1 specific antibodies, followed by protein identification using mass spectrometry (IP-MS). As control, we precipitated PID-1 from two independent *pid-1* loss of function strains. Both experiments identified a rather restricted set of proteins (Figure 1a, S1a). TOFU-6 and PID-3 were identified as the most prominent PID-1 interactors, and for the latter we further validated the interaction with PID-1 through IP-Western blotting (Figure S1b). Given that these two proteins were identified in an RNAi-screen for 21U biogenesis factors ^27^, we considered these as functionally relevant PID-1 interactors. TOFU-6 is a protein with a Tudor domain, a potential eIF4E interaction motif and an RRM domain, whereas PID-3 has an RRM domain and a MID-domain (Figure S1c). Two other proteins were consistently identified as PID-1 interactors: IFE-3 and ERH-2 (Figure 1a, S1a). IFE-3 is one of the five *C. elegans* homologs of eIF4E. Previous work demonstrated that IFE-3 binds to the 7-methylguanylate (m7G) Cap ^34^. Interestingly, 21U precursor transcripts appear to be not trans-spliced ^25^, implying that they have a m7G Cap structure at their 5’ end. Hence, IFE-3 may bind to the 5’ cap structure of the 21U precursor transcripts. ERH-2 is one of the two *C. elegans* homologs of a protein known as ‘enhancer of rudimentary’. Homologs of this protein are strongly conserved from plants to mammals. Even though a protein structure for the human homolog has been described ^35^, its molecular function is still unclear. In Figure S1c we present a schematic of all these PID-1 interactors, with their identified domains.

**Figure 1.**
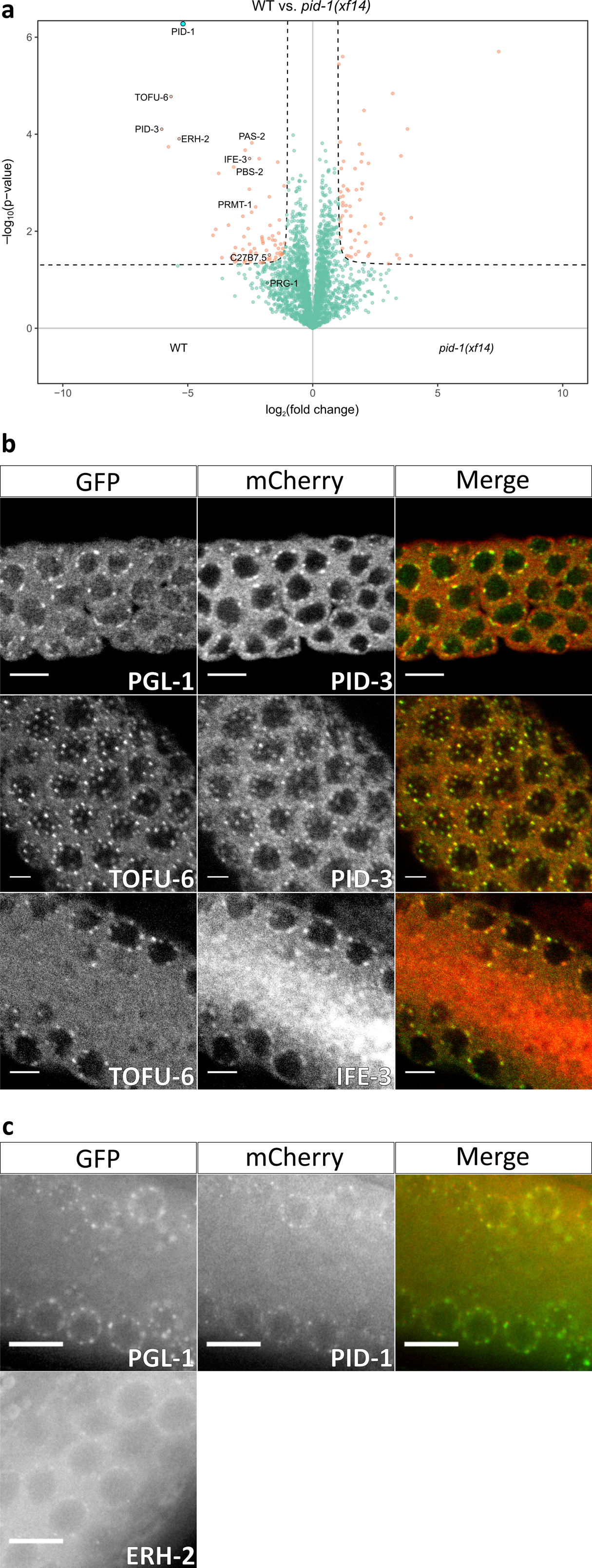
PID-1 interactors reside in P-Granules. **a)** Volcano plot representing label-free proteomic quantification of PID-1 IPs from non-gravid adult extracts. IPs were performed and analyzed in quadruplicates. The x-axis represents the median fold enrichment of individual proteins in wild type (WT) versus *pid-1(xf14)* mutant strain. y-axis indicates -Log_10_(p-value) of observed enrichments. Dashed lines represent thresholds at p=0.05 and 2-fold enrichment. Blue data points represent values out of scale. Red and Green data points represent values above and below threshold, respectively. **b and c)** Expression pattern and localization of tagged PETISCO components and P-granule marker PGL-1. Endogenous promotors and 3’UTRs were used for PETISCO proteins. Proteins and observable tags indicated in the panels. **b)** Immunostaining images acquired with laser scanning confocal microscope. **c)** Live worm images acquired under wide field fluorescent microscope. Scale bars represent 5 🕴m. Contrast of images has been enhanced.

We next probed the subcellular localization of PID-1 and its four identified interactors, by expressing them from transgenes generated through the so-called miniMos approach (see methods) ^36^. For each gene we used its own promoter and 3’ UTR sequences. All transgenes were able to rescue mutant phenotypes (not shown) indicating the expressed proteins are functional. All four interactors show a characteristic perinuclear, punctate localization in the germline syncytium, overlapping partially, but not fully with the P-granule marker PGL-1 (Figure 1b,c). IFE-3 has a clear granular expression in the primordial germ cells in the embryo (Figure S2b), whereas the remaining interactors show a dispersed cytoplasmic distribution across the entire early embryo (Figure S2c). These results show that all the identified PID-1 interactors are expressed concomitantly in early embryos, and in the germline, where they are found in close proximity to P-granules.

### PID-1 interactors mutually interact to form PETISCO

To probe to what extent the identified PID-1 interactors reciprocally interact, and to what extent they in turn interact with additional proteins, we performed IP-MS on TOFU-6, IFE-3, PID-3 and ERH-2 in young adult animals, using the tagged proteins expressed from the above-described transgenes. Extracts from non-transgenic wild-type animals were used as negative controls. As shown in Figure 2a-d, these experiments revealed that all four proteins co-precipitate with each other. In addition, we also found an uncharacterised protein (C35D10.13) which systematically co-precipitated with the PID-1 interactors, but was absent from the PID-1 IPs. This factor, which we named TOST-1, will be further described below. These interactions are RNAse resistant. RNase treatment did not disrupt PID-3 interactions (Figure S3a), and had very little effect on IFE-3 partners, resulting only in the loss of PID-1 (Figure S3b). We summarize the network of PID-1 interactors in Figure 2e. Given the strong reciprocal interactions between these proteins, it is likely that they form a discrete complex. We named this complex PETISCO, for <u>P</u>ID-3, <u>E</u>RH-2, <u>T</u>OFU-6, IFE-3 <u>s</u>mall RNA <u>Co</u>mplex.

**Figure 2.**
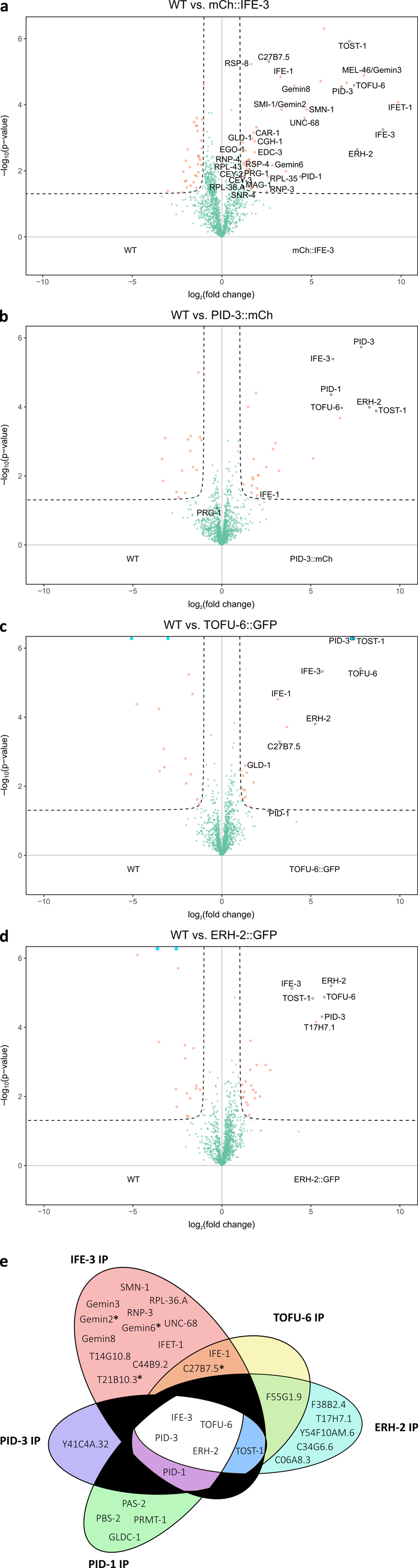
PID-1 interactors form a novel protein complex, PETISCO. **a-d)** Volcano plot representing label-free proteomic quantification of quadruplicate **a)** 3xFLAG::mCherry::IFE-3;*ife-3(xf101)*; **b)** PID-3::mCherry::Myc;*pid-3(tm2417)*; **c)** TOFU-6::GFP::HA;*tofu-6(it20)* and **d)** ERH-2::GFP::OLLAS;*erh-2(xf168)* IPs from non-gravid adult extracts. The x-axis represents the median fold enrichment of individual proteins in control (WT) versus transgenic strain. y-axis indicates -Log_10_(p-value) of observed enrichments. Dashed lines represent thresholds at p=0.05 and 2-fold enrichment. Blue data points represent values out of scale. Red and Green data points represent above and below threshold respectively. **e)** Venn Diagram summarizing significant interactions in PETISCO protein IPs. *represents protein found significantly enriched in only one experiment of 3xFLAG::mCherry::IFE-3;*ife-3(xf101)* IP.

As mentioned before, IFE-3 is one of the *C. elegans* eIF4E homologs. Surprisingly, in our experiments we do not detect any of the known translation initiation factors to be associated with IFE-3. The transgenically expressed IFE-3 does rescue the *ife-3* mutant phenotype (not shown), implying that IFE-3 may not play a role in initiating translation. We do detect a number of additional proteins bound to IFE-3 which provide clues to IFE-3 function. One such protein is IFET-1, a homolog of human EIF4E nuclear import factor 1 and a negative regulator of translation ^37^. We further consistently detect many, if not all components of the SMN complex ^38^. The SMN complex plays a major role in snRNP biogenesis, implicating IFE-3 in this process as well.

### PETISCO architecture

Having established the components of PETISCO, we next dissected the molecular interactions within this complex. Using the yeast-two-hybrid (Y2H) system we scored interactions between each possible pair of individual PETISCO subunits (Figure 3a, S4a). This resulted in the following interactions that are most likely to be direct: IFE-3 interacts only with TOFU-6. TOFU-6 in turn interacts with PID-3, which in turn also interacts with ERH-2. Finally, PID-1 and TOST-1 were both found to interact only with ERH-2. Self-interaction was detected for ERH-2, though under lower stringency conditions, indicating that ERH-2 may be present as a dimer in PETISCO.

**Figure 3.**
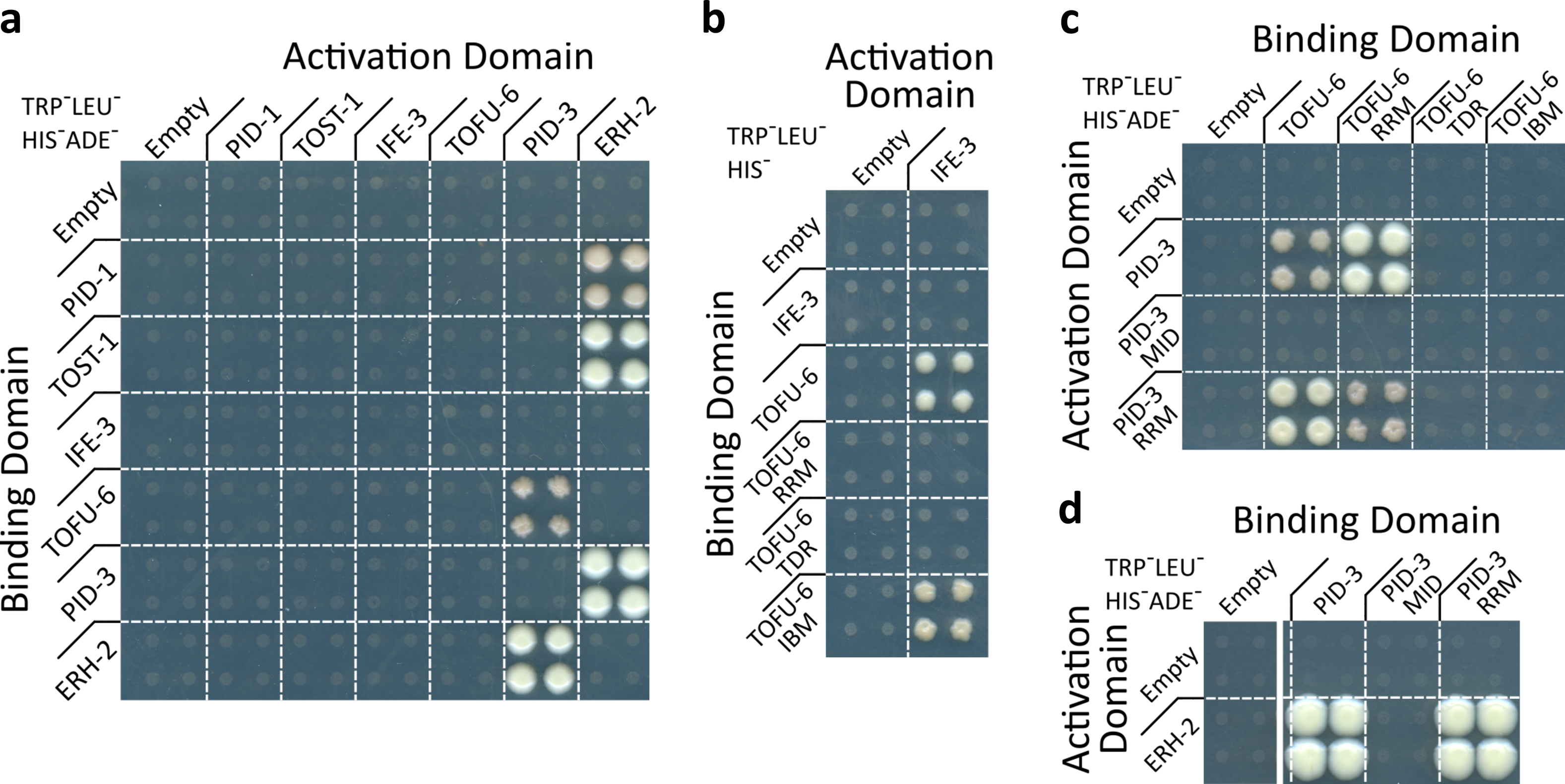
PETISCO Architecture. **a-d)** Yeast two-hybrid interaction assays of PETISCO subunits in low (TRP^-^LEU^-^HIS^-^) or high stringency media (TRP^-^LEU^-^HIS^-^ADE^-^) as indicated. Interactions were screened in both Y2H orientations. **a)** Full length proteins **b)** TOFU-6 and individual domains tested for interaction with full length IFE-3 **c)** Interactions between PID-3 and TOFU-6 **d)** Interaction between PID-3 and ERH-1. For details on domains and other selection conditions see Figure S4.

We also investigated which domains of the PETISCO subunits are responsible for the protein interactions using the same Y2H set-up. The putative eIF4E interaction motif ^39 40^ within the C-terminus of TOFU-6 indeed is responsible for the interaction between TOFU-6 and IFE-3 (Figure 3b). No interactions were found for the Tudor domain of TOFU-6. The RRM domain of TOFU-6 was found to bind to the RRM-domain of PID-3 (Figure 3c), and the same RRM domain of PID-3 was found to also interact with ERH-2 (Figure 3d). Lower stringency and control selections of the same experiments are shown in Figures S4b-d). This Y2H set-up did not allow us to determine if the PID-3 RRM domain can sustain both the TOFU-6 and the ERH-2 interaction simultaneously, however the fact that we find PID-3, TOFU-6 and ERH-2 strongly enriched in the IPs of one another, suggests that it can.

### PETISCO subunits are required for 21U RNA biogenesis and are essential for embryogenesis

Since at least two components of PETISCO, PID-1 and TOFU-6, play a major role in 21U RNA biogenesis, we hypothesized that the other components are also part of this pathway. We first tested whether they affect the silencing of a GFP sensor construct that reports on the activity of the 21U pathway (21U sensor) ^11^. We used a strain that contains this sensor plus a *pid-1(xf14)* mutation that is rescued by transgenic expression of PID-1. In this strain, the sensor is silenced (Figure 4a,b), although not completely. This partial 21U sensor-silencing brings two advantages: first, the 21U sensor is not in an RNAe-state, in which it would no longer be activated by loss of 21U RNAs; second, a semi-silenced state allows for scoring of both increased and decreased sensor activity.

**Figure 4.**
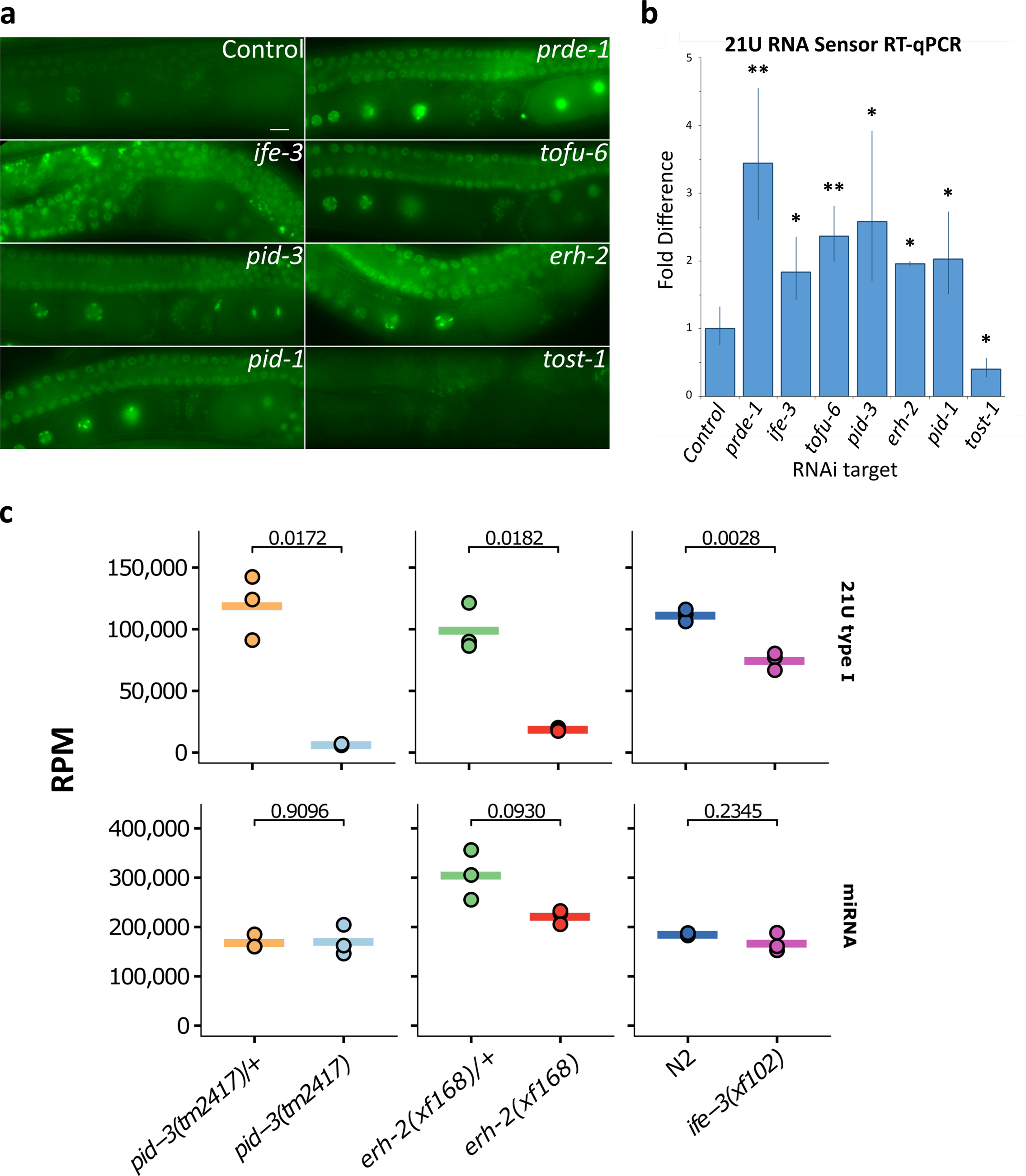
PETISCO is required for 21U RNA biogenesis. **a)** Wide field fluorescent microscopy of adult hermaphrodites carrying GFPH2B-21U sensor transgene in a sensitized background. Worms were subjected to RNAi via feeding (targets indicated in figure) from L1 larval stage to adulthood. Empty RNAi Vector serves as negative control. Scale Bar 10 🕴m. **b)** Quantitative RT-PCR of 21U sensor transgene in adult worm populations of A). Values are obtained from experimental triplicates and technical duplicates, normalized to *pmp-3* mRNA levels. Significance was tested with Student’s t-test: **p-value<0.01; *p-value<0.05. Error bars represent standard deviation. **c)** Global level of type I 21U RNA and miRNAs in the indicated strains. Values are in reads per million (RPM). Individual data points of three independent replicates are shown and horizontal bar represents the mean. Significance was tested with Student’s t-test and p-values are indicated in the graph.

Animals from this sensitized sensor strain were subjected to RNAi targeting the various PETISCO subunits and sensor activity was evaluated by microscopy and RT-qPCR (Figure 4a,b). As expected, RNAi against *pid-1* activated the sensor. Likewise, RNAi against *ife-3*, *pid-3*, *tofu-6* and *erh-2* activated the sensor. These data show that PETISCO subunits, like the PETISCO interactor PID-1, are required for a fully functional 21U RNA silencing pathway.

To extend the above observation to the overall 21U RNA population, we aimed to sequence small RNAs from genetic mutants. For this, we created mutant alleles using CRISPR-Cas9 for each of the PETISCO subunit genes. We were able to derive deletion alleles for *pid-3*, *erh-2* and *ife-3* (Figure S5a), and isolated and sequenced small RNAs. We did not make mutants for *tofu-6*, since RNAi on *tofu-6* was already shown to significantly reduce 21U RNA production ^27^. All experiments were done in triplicates. The results show that all tested PETISCO subunits significantly affect 21U RNA biogenesis (Figure 4c). In particular, *pid-3(tm2417)* and *erh-2(xf168)* mutants display a strong reduction in 21U RNA levels. The effect of *ife-3(xf102)* mutation on 21U RNA accumulation is less pronounced, but still significant. Redundancy between IFE-3 and other eIF4E homologs, could be a reason for this less pronounced effect on 21U RNA levels, as we find IFE-1, a non-selective TMG/m7G binder ^34^, enriched in some of our IP-MS experiments (Figure 2a-c, S3b). Other types of small RNAs, such as 22G, 26G and miRNAs were mostly not affected by the tested mutations (Figure 4c and S5b-d). We note that so-called type II 21U RNAs, which come from loci lacking the canonical Ruby motif and are expressed at much lower levels ^25^, are only mildly, or not at all affected by loss of PETISCO components.

Besides the defects in 21U biogenesis, the mutant alleles also display a so-called maternal effect lethal (Mel) phenotype: homozygous mutant offspring from a heterozygous animal develop into fertile adults, but the embryos display a fully penetrant arrest and never hatch (Table 1). This phenotype has already been described for *tofu-6*, which is also known as *mel-47* ^41^. The *ife-3* mutant displays a mixed phenotype, as previously described ^42^: homozygous mutant offspring from a heterozygous animal develop into adults that can either display a masculinized germline (Mog) or a Mel phenotype. Interestingly, we note that *pid-1* mutants also display a Mog phenotype, albeit at low frequency (Table 1 and Figure S6).

**Table 1.** PETISCO displays maternal effect lethality. “n.a.” not applicable; “n.d.” not determined; “-“ mild 21U RNA defect; “--“ severe 21U RNA defect; “+” no 21U RNA defect; “(TS)” temperature sensitive; “#” counts are for gonadal arms (n=38) due to mixed phenotypes in individuals; “*” according to Goh *et al*. ^27^.

| Gene | Allele | Mutation Type | 21U RNA Presence | Terminal Phenotypes | Ratio (%) |
| --- | --- | --- | --- | --- | --- |
| *ife-3* | *xf101* | Start Codon Loss | n.d. | Mog/Mel | n.d. |
|  | *xf102* | Frameshift | - | Mog/Mel | n.d. |
|  | RNAi | n.a. | n.d. | Mog/Mel | 47/53# |
| *pid-3* | *tm2417* | Frameshift | -- | Mel | 100 |
|  | *xf149* | Frameshift | n.d. | Mel | 100 |
|  | *xf151* | Inframe Deletion | n.d. | Mel | 100 |
|  | *xf153* | Frameshift | n.d. | Mel | 100 |
| *tofu-6* | *it20* | Nonsense | n.d. | Mel | 100 |
|  | *yt2* | Nonsense | n.d. | Mel | 100 |
|  | RNAi | n.a. | --* | Mel | n.d. |
| *tost-1* | *xf191* | Frameshift | n.d. | Viable | n.d. |
|  | *xf194* | Start Codon Loss | + | Mel | 100 |
|  | *xf196* | Splice Site Loss | n.d. | (TS) Mel | 100 (25°C) |
| *pid-1* | *xf14* | Missense | -- | n.d. | n.d. |
|  | *xf35* | Frameshift | -- | Mog | <1 |
|  | *xf36* | Frameshift | -- | n.d. | n.d. |
| *erh-2* | *xf168* | Stop Codon Loss | -- | Mel | 100 |

### PID-1 and TOST-1 define distinct functions of PETISCO

The fact that PETISCO genes are essential for embryogenesis contrasts with the fact that loss of 21U RNAs through other means, such as loss of PRG-1 or PID-1, does not result in a Mel phenotype. Interestingly, RNAi-knockdown of *tost-1* resulted in an embryonic lethal phenotype (not shown). In our IP-MS experiments, we consistently found this protein to interact with the PETISCO subunits (Figure 2a-d), yet not with PID-1 (Figure 1a and S1a). Furthermore, *tost-1(rnai)* displays enhanced, rather than disrupted silencing of the sensitized 21U sensor (Figure 4a,b). This phenotype lead us to name this protein TOST-1, short for <u>t</u>wenty-<u>o</u>ne U antagoni<u>st.</u>

We tested both PID-1 and TOST-1 for interactions with PETISCO subunits. We found that both PID-1 and TOST-1 specifically interact only with ERH-2 (Figure 3a). While the overall amino-acid sequences of PID-1 and TOST-1 do not display convincing homology (Figure S7a), when aligned with PID-1 and TOST-1 homologs from other nematode species, a conserved motif emerges (_[+][+]Ψ(T/S)_**L**_(N/S)[-]_**RF**_xΨxxx_**G**_(Y/F)_ – Figure 5a). Strikingly, the *xf14* allele we identified for *pid-1* in our previously described genetic screen caries a mutation of the fully conserved arginine residue within this motif (R61C) ^26^. When introduced into the Y2H experiment, PID-1(R61C) did not interact with ERH-2, and the analogous mutation in TOST-1 also disrupted its ERH-2 interaction (Figure 5b). These data strongly suggest that both PID-1 and TOST-1 share a conserved short motif required for ERH-2 interaction.

**Figure 5.**
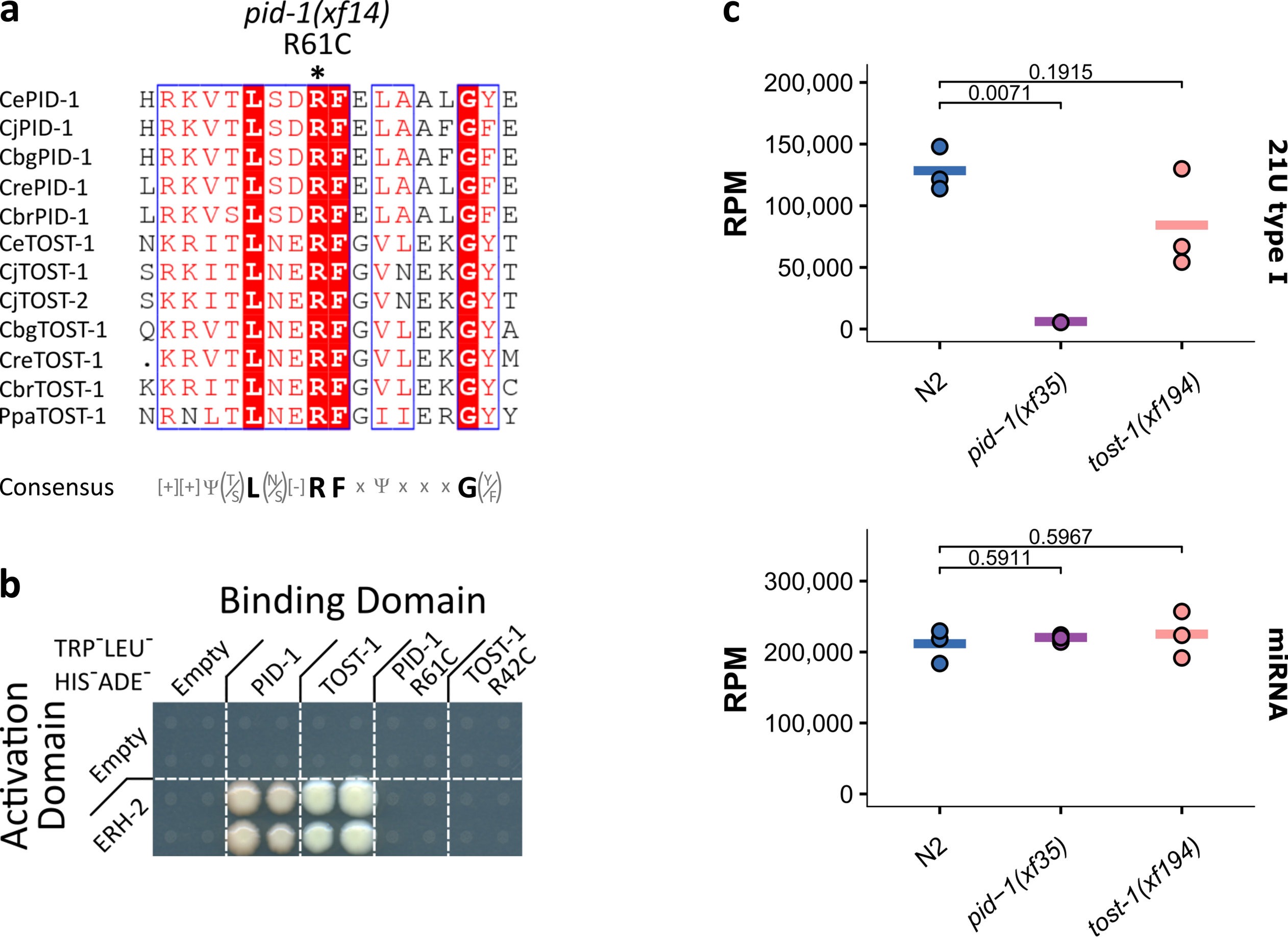
TOST-1 is essential but not required for 21U RNA biogenesis. **a)** Alignment of a short region of PID-1 and TOST-1 homologs from different nematodes. Conserved arginine residue was found to be mutated in *pid-1(xf14)* as indicated. Consensus sequence is presented below alignment. Alignment performed with MUSCLE v3.8 ^50^ and representation with ESPrit v3.0 ^51^. **b)** Y2H interaction assay of PID-1, TOST-1 and PID-1/TOST-1 carrying the corresponding arginine mutation found in pid-1(xf14). High stringency plates (TRP^-^LEU^-^HIS^-^ADE^-^) were used in the presented figure. For other conditions please see Figure S4d. **c)** Global levels of type I 21U RNA and miRNAs in wild type (N2), *pid-1(xf35)* and *tost-1(xf194)* gravid adult worms. Values are in reads per million (RPM). Individual data points of three independent replicates are shown and horizontal bar represents the mean. Significance was tested with Student’s t-test and p-values are indicated in the graph.

To further understand TOST-1 function we generated loss of function alleles for *tost-1* using CRISPR-Cas9 (Figure S5a). Loss of TOST-1 did not significantly affect 21U RNA biogenesis (Figure 5c, S7b), but did result in a fully penetrant Mel phenotype (Table 1). Among the alleles we generated, we found one allele, *tost-1(xf196)*, with a 58bp deletion which removes the splice acceptor site of its third exon (Figure S5a), resulting in either a frame-shift or a C-terminal truncation. *tost-1(xf196)* displays a temperature-sensitive (TS) effect: at 25°C the animals are fully Mel, while at 15°C the animals are viable. The fact that this allele appears to be mostly functional at 15°C suggests that the N-terminal part of TOST-1 is the critical part of this protein. The interaction motif we identified is still intact in *tost-1(xf196),* consistent with the idea that it is essential for TOST-1 function. This TS allele allowed us to probe the Mel phenotype in more detail, through temperature-shift experiments (Figure S7c). First, animals grown at the restrictive temperature were able to produce viable offspring after shifting them to the permissive temperature. This shows that there are no structural or developmental defects in the germline of these animals that prevent them from producing live offspring. Second, the TS allele allows us to probe when TOST-1 function is required. The time required for animals shifted to the permissive temperature to produce viable embryos (8 hours) is significantly longer than the time it takes from fertilization to egg-laying (approximately 200 min at 15°C), implying that the embryos laid after 8 hours were fertilized and raised at the permissive temperature. Yet, they are still arresting. This observation places TOST-1 activity within the gonad. However, when animals are shifted from permissive to restrictive temperature, the first arrested embryos are laid already after two hours. This is very close to the time of residency within the uterus (approximately 150 min 25°C), and hence these embryos had been fertilized very close to the time of the temperature shift, in a gonad that was still functional. This suggests TOST-1 also has a function within the embryo. This is consistent with PETISCO subunit expression within early embryos (Figure S2c).

In conclusion, our data shows that both TOST-1 and PID-1 bind to the same PETISCO subunit, while loss of these two proteins results in different phenotypes. The combined phenotypes of both *pid-1* and *tost-1* mutants is found in mutants for the other PETISCO subunits, strongly suggesting that PID-1 and TOST-1 define two distinct aspects of PETISCO function.

### PETISCO interacts with SL1 RNA

Given the known links between the 21U RNA pathway and snRNA transcription, the fact that we find many SMN-complex components in IFE-3 IPs, and that at least one SMN component has a Mel phenotype (MEL-46), we hypothesized that the essential function of PETISCO is linked to snRNA homeostasis. Consistent with this idea, we noticed a striking accumulation of a 3’ fragment of SL1 RNA in a number of the mutants we tested (Figure 6a,b, S8a). Such accumulation was less pronounced, or absent for another small non-coding RNA that is produced from the same gene cluster as SL1, 5S rRNA (Figure 6b,c). Importantly, this enrichment of SL1 fragments was very clear in *tost-1* mutants, but absent in *pid-1* mutants. Also *ife-3* mutants and *erh-2* mutants displayed a similar accumulation in all the replicates that we sequenced. Mutants for *pid-3*, however, did not show the accumulation (Figure 6b). For SL2 we only observe the enrichment in *ife-3* mutant libraries (Figure S8b), but we point out that SL2 read counts are comparatively very low.

**Figure 6.**
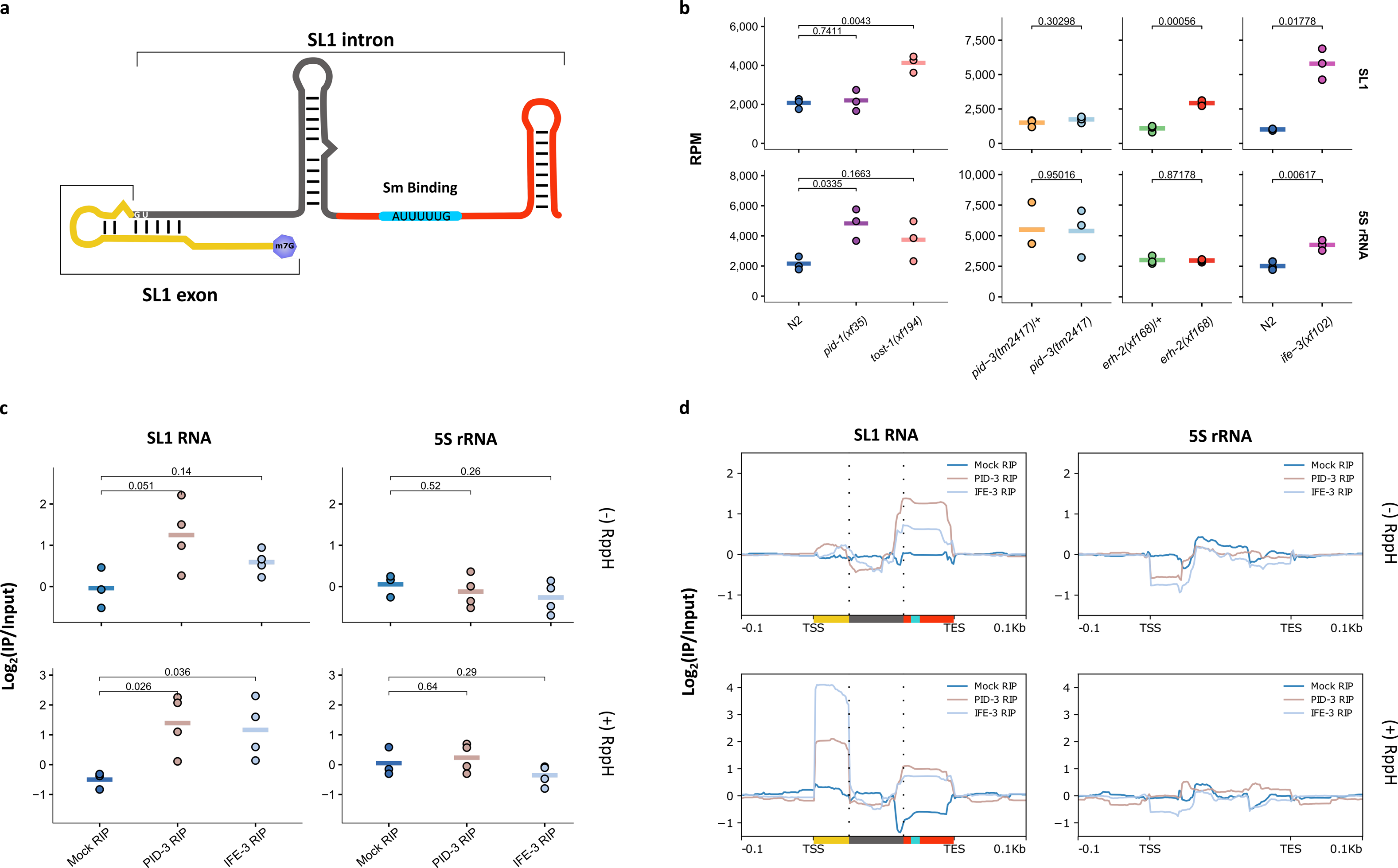
PETISCO interacts with SL1 snRNA. **a)** Schematic representation of the Splicing Leader 1 RNA. **b)** Global levels of SL1 RNA and 5S rRNA in wild type (N2), *pid-1(xf35)* and *tost-1(xf194)* gravid adult worms and *pid-3(tm2417), erh-2(xf168)* and *ife-3(xf102)* non-gravid adults. Values are in reads per million (RPM). Individual data points of three independent replicates are shown and horizontal bar represents the total mean. Significance was tested with Student’s t-test and p-values are indicated in the graph. **c)** Fold enrichments of SL1 RNA and 5S rRNA in Mock (N2), 3xFLAG::mCherry::IFE-3;*ife-3(xf101)*; and PID-3::mCherry::Myc;*pid-3(tm2417)* RIPs over paired inputs in non-gravid adult worms. Top row displays non-treated and bottom row RppH treated samples. Individual data points of three independent replicates are shown and horizontal bar represents the mean. Significance was tested with Student’s t-test and p-values are indicated in the graph. **d)** Coverage profile, normalized to paired input, of SL1, of the data displayed in **c**. Colors under SL1 RNA correspond to scaled colors represented in **a.**

**Figure 7.**
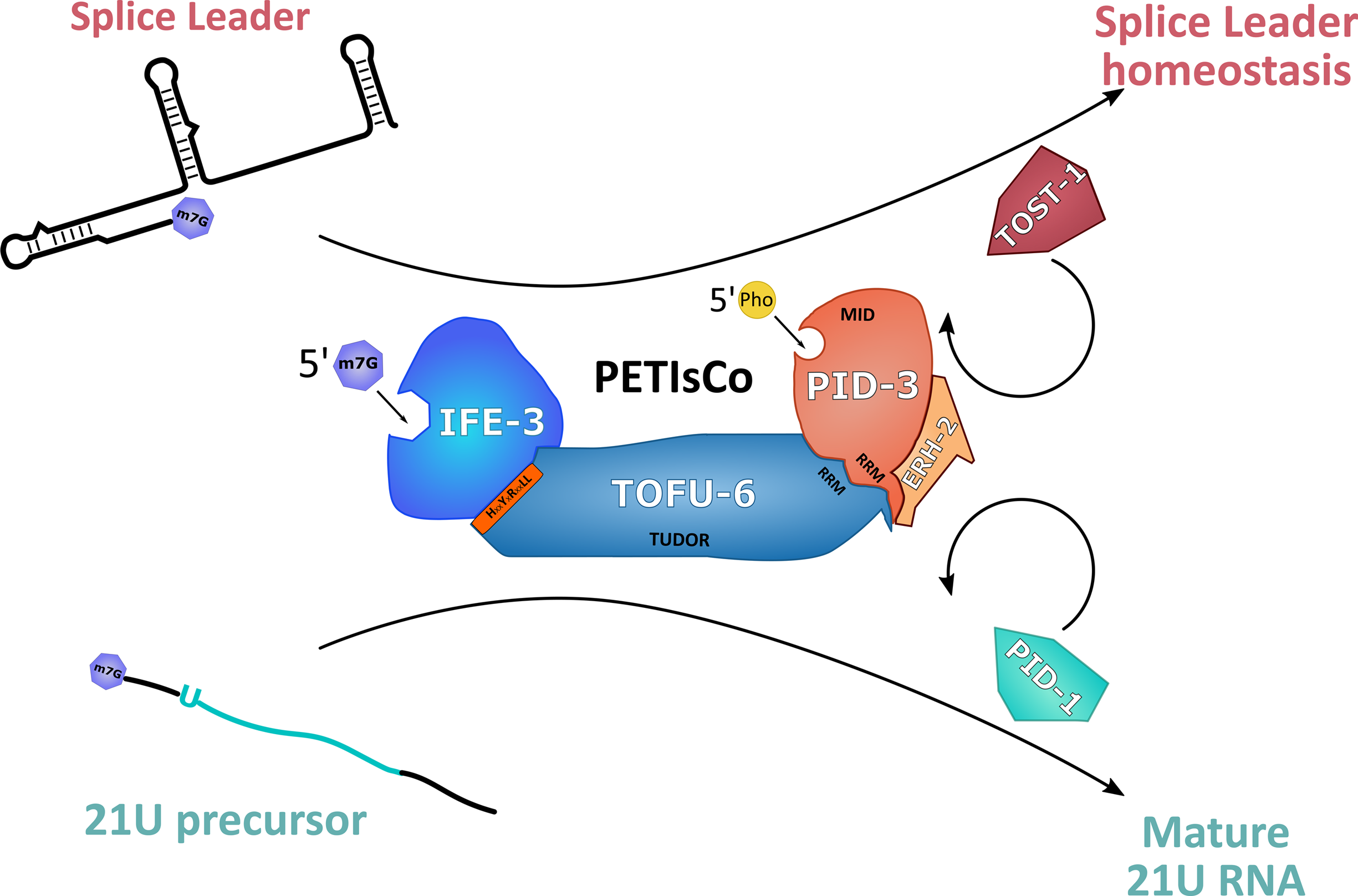
Schematic representation of PETISCO function. A schematic of the proposed PETISCO activity, displaying its dual function. One in 21U biogenesis and another potentially linking to splice leader homeostasis. The two different 5’ end binding domains may reflect stabilization of different RNA species, and may reflect 5’ end processing of transcripts bound by PETISCO. ERH-2 serves as an anchor for PID-1 or TOST-1 driving PETISCO function towards 21U RNA biogenesis or SL1 homeostasis respectively.

To further test a potential link between PETISCO and SL1 RNA, we probed whether SL1, or SL1 fragments may be bound by PETISCO. To test this, we performed RIPseq experiments on IFE-3 and PID-3. We performed these experiments in quadruplicate, and also performed triplicate mock IPs from wild-type animals. Since full SL1 transcripts, but also 21U precursors are capped, we treated the RNA with RppH to remove 5’ caps before cloning. We detect strong and significant enrichment of SL1, and to a lesser extent SL2 sequences in both PID-3 and IFE-3 IPs (Figure 6c, S8c). In contrast, transcripts from 21U loci, 5S rRNA and snRNAs are not enriched, or are even depleted (Figure 6c, S8c). The read coverage over the SL1 transcript shows that two fragments are detected: one at the 5’ and another at the 3’ end (Figure 6d, S8d). The 5’ fragment is capped, as this fragment is absent from libraries from non-RppH treated RNA (Figure 6d). We conclude that PETISCO binds to SL1 RNA and that loss of PETISCO leads to an accumulation of SL1 RNA fragments.

## Discussion

We have identified a protein complex, named PETISCO, that is involved in at least two different pathways. PID-1 guides PETISCO towards 21U RNA biogenesis, whereas TOST-1 brings PETISCO to a pathway that is essential for embryogenesis, but does not involve 21U RNAs. Here we discuss various aspects of this complex and present hypotheses for the different potential functions of PETISCO.

### PID-3 and TOFU-6 define the core of PETISCO

Judging by the enrichments found in all IPs and the similarities in phenotypes of the corresponding mutants it seems likely that two proteins, PID-3 and TOFU-6 form the core of the PETISCO complex, and that the other identified factors may add as adaptors for specific functions of PETISCO. What could the core function(s) of this complex be? Based on protein identities and obtained results we consider the following, non-exclusive possibilities.

Based on the presence of two RRM domains, one in TOFU-6 and one in PID-3, this complex likely has RNA binding activities. Indeed, our RIPseq experiment identified 3’ fragments of SL1 and SL2 bound to PETISCO. Interestingly, we found that both RRM domains act as protein-protein interaction modules. This does not mean that these two RRM domains cannot also be involved in RNA binding, as combined RNA- and protein-binding activities for RRM domains have been described before ^43^. Possibly, the assembly of the RRM-RRM contact between PID-3 and TOFU-6 is facilitated by RNA. Even though we find that RNase treatment does not affect the pull-down efficiency between PID-3 and TOFU-6, the RNA involved may be inaccessible to RNases.

In addition to the RRM motifs, the MID-domain present in PID-3 also brings potential for protein-RNA-interactions. The MID-domain has thus far been described in context of argonaute proteins where it mediates interactions with the 5’ end of the bound small RNA co-factor ^44^. It is curious to find such a domain in a protein involved in generating small RNA co-factors for an argonaute protein. We hypothesize that the MID domain of PID-3 may help to stabilize a 5’ processing intermediate. This would imply PETISCO as a 5’end processing platform for 21U RNAs. Biochemical reconstitution experiments will be required to test this.

The Tudor domain of TOFU-6 likely represents a protein-protein interaction unit. Tudor domains are known to interact with methylated arginines, and within perinuclear granules, where PETISCO is found, such methylated arginines are likely abundant ^45^. Hence, the Tudor domain may be involved in sequestering PETISCO to these granules, possibly to PRG-1. PRG-1 itself has an N-terminal protein sequence that is highly suggestive of arginine methylation. In one of our IFE-3 pull-downs, we do enrich for PRG-1 (Figure 2a), supporting this possibility.

### ERH-2: Bridge between core-PETISCO and PETISCO-guiding factors?

The presence of ERH-2 provides hints for PETISCO function. In *S. pombe*, Erh1 forms a tight complex with a protein named Mmi and together they are involved in nuclear mRNA degradation of meiotic transcripts, involving proteins such as ARS2 and the CCR4-NOT complex ^33^. Hence, Erh1 in *S. pombe* appears to act as a bridge between an RNA binding protein (Mmi) and an RNA processing machinery. Our data strongly suggest that ERH-2 in *C. elegans* has a very similar function. On the one hand, it binds core PETISCO and on the other hand either PID-1 or TOST-1, which set the function of the complex (see below).

ERH-related proteins in other systems, including *S. pombe* and human cells, are nuclear proteins ^33^, while we find ERH-2 to be present mainly in perinuclear granules. Related to this it is interesting to note that *C. elegans* has a second ERH-like protein T21C9.4. Alignments show that this homolog is more closely related to *S. pombe* and human ERH1 (not shown), raising the possibility that T21C9.4 in *C. elegans* is nuclear and may be coupled to nuclear RNA processing.

### Bi-functionalization of PETISCO through PID-1 and TOST-1

We show that PETISCO has at least two functions: 21U RNA biogenesis and another, essential function in embryogenesis, which is likely related to SL1/2 snRNP homeostasis. These two functions can be functionally separated through two different ‘adapter’-proteins (PID-1 and TOST-1) that bind to PETISCO via ERH-2. The overall homology between PID-1 and TOST-1 is very low. Nevertheless, we identified a motif that is required for binding to ERH-2. Hence, both proteins likely bind to PETISCO in a mutually exclusive way. Indeed, no TOST-1 was detected in PID-1 IPs, and Zeng et al. (accompanying manuscript) did not find PID-1 in their TOST-1 IPs. An IP-MS experiment on PID-3 in absence of PID-1 (therefore enriching for TOST-1:PETISCO) did not reveal much change compared to wild-type (not shown), suggesting that stable interactions within PID-1:PETISCO and TOST-1:PETISCO are very similar and do not provide further insights into the differential activities of PID-1 and TOST-1.

### Potential roles for IFE-3

IFE-3 has been shown to bind m7G Cap structures, and to bind much less efficiently to the typical TMG Cap structures found on the majority of trans-spliced mRNAs in *C. elegans* ^34^. This activity of IFE-3 makes it a good candidate for binding 21U RNA precursors at a stage before 5’ processing. IFE-3 IP-MS experiments also enrich mildly for proteins involved in de-capping (e.g. EDC-3, CAR-1, CGH-1 – Figure 2a). In relation to 21U RNA biogenesis, it is an intriguing option that IFE-3 binds 21U precursor transcripts through their 5’ cap structure, after which they become de-capped by associated de-capping activities. IFE-3, however, is not essential for 21U generation. This could be due to either functional redundancy with IFE-1, which we regularly detect in our IP-MS experiments. Alternatively, IFE-3 has no direct role in 21U RNA processing, and the small drop of 21U levels in *ife-3* mutants is an indirect effect, for instance through effects on PETISCO overall stability.

We find that IFE-3 consistently comes down with many, if not all proteins found in the so-called SMN complex. This complex is well known for its involvement in snRNP assembly ^38^. In *C. elegans*, the SMN complex has also been shown to be required for SL1 trans-splicing ^46^. We note that a homolog for Gemin5, which has been shown to bind the m7G Cap structure of pre-mature snRNAs in human cells ^47^, is not encoded by the *C. elegans* genome. Given the cap-binding activity of IFE-3 and its association with the other SMN complex subunits, IFE-3 may fulfil this function in *C. elegans*. We note that like loss of IFE-3 ^42^, loss of the *C. elegans* Gemin3 homolog, named MEL-46 ^41^, and the U2 snRNP-associated factor MOG-2 ^48^ also result in Maternal effect lethality (Mel) and Masculinization of the germline (Mog) phenotypes, further strengthening the links between IFE-3 and snRNP homeostasis. Given that the SMN-complex proteins are not found in the IPs of any of the other PETISCO subunits, the IFE-3-SMN interaction is likely to be physically separated from PETISCO. Finally, it is interesting to note that *pid-1* mutants, but not *tost-1* mutants, also display a Mog phenotype, albeit at a low frequency. Considering the interplay between PID-1 and TOST-1, this could relate from excessive, or ectopic TOST-1:PETISCO in *pid-1* mutants affecting the IFE-3-SMN interplay.

### Potential mechanisms behind the Mel and 21U phenotypes of PETISCO mutants

Our data clearly show that PETISCO has at least two functions. Its role in 21U RNA biogenesis is non-essential and is guided by PID-1. A second function, which is essential for early development, is linked to PETISCO via TOST-1. While our current data do not provide mechanistic details on molecular PETISCO function, they do provide interesting leads as to what the essential function of PETISCO may be.

As mentioned above, PETISCO contains proteins with domains that interact with the 5’ ends of RNA, either capped or phosphorylated, raising the possibility PID-1-bound PETISCO plays a role in 5’ end processing of 21U precursor transcripts. In our RIP experiments, we did not detect 21U RNA precursor molecules, suggesting that either levels are too low to be detected in a RIP experiment, or that their processing is too fast. Given that RIP experiments typically have high backgrounds, this may easily prevent detection of significant enrichments. In view of the phenotypes and protein domains within PETISCO, we consider it very likely that PETISCO does interact with 21U RNA precursors.

But what could the molecular function of PID-1-bound PETISCO be? In absence of PETISCO 21U RNA levels drop strongly, but we note that so-called type II 21U RNA levels are much less affected (Figure S5b). These 21U RNA species do not derive from loci with the typical Ruby-motif, and are expressed at much lower levels than type I 21U RNAs. We hypothesize that PETISCO may function to specifically stimulate the processing of pre-cursors that come from Ruby-motif-containing 21U loci, such that the PRG-1 RNP pool is dominated by PRG-1 protein bound to type I 21U RNAs.

TOST-1-bound PETISCO may have a similar function, only then coupled to the production of an RNP that is essential for early development. Our RIPseq and mutant small RNAseq data indicate that this could involve the SL1 snRNP. Our interpretation of these data is that PETISCO binds full-length SL1 transcripts, but that these are partially degraded by nucleases during the RIP-procedure, or within the animals when further processing of the complex is stalled due to loss of PETISCO subunits such as TOST-1 or IFE-3. But how would this explain the maternal effect lethality phenotype of PETISCO mutants? SL1 is a generally required splice-leader RNA, and mutants that carry a large deletion covering the SL1 loci display a direct embryonic lethal phenotype ^49^. If PETISCO would have a core function in trans-splicing one would expect a similarly severe zygotic phenotype. We hypothesize that PETISCO specifically helps the accumulation of a large pool of SL1 snRNPs that will be maternally loaded into the embryos. Later in development, when the demand for SL1 snRNPs may have dropped, PETISCO may therefore not be required. This would create an interesting parallel to 21U RNPs, that are also provided maternally ^15^.

Clearly, additional studies will be required to resolve the molecular function of PETISCO in 21U RNA processing and further test its role in SL1 homeostasis during early development. However, our data firmly show that PETISCO is a 21U biogenesis complex, and in addition has an essential role during early development. Our work shows that the processing of 21U RNAs is likely derived from a much more widely conserved mechanism, and provides a new and exciting view into how small RNA biogenesis can be intertwined with other gene-regulatory mechanisms.

## Acknowledgements

We thank all the members of the Ulrich and Ketting lab for great help and discussion. A special thanks to the Yasmin El Sherif and Sonja Braun for technical assistance. We thank Miguel Andrade, Peter Sarkies and Ian MacRae for helpful discussions. The authors are grateful to Hanna Lukas, Clara Werner and Maria Mendez-Lago of the IMB genomics core facility for library preparation. We thank the IMB Media Lab, Microscopy and Cytometry Core Facilities for consumables and equipment. Some strains were provided by the Caenorhabditis Genetics Center (CGC), which is funded by NIH Office of Research Infrastructure Programs (P40 OD010440) and the National BioResource Project, managed by the Mitani Lab. This work was supported by Deutsche Forschungsgemeinschaft grant KE 1888/1-1 and KE1888/1-2 (Project Funding Programme to R.F.K.), ERC-StG 202819 (RFK) and ERC-AdG 323179 (HDU).

## References

1. Ketting, R.F. The many faces of RNAi. Dev Cell 20, 148-61 (2011).

2. Weick, E.M. & Miska, E.A. piRNAs: from biogenesis to function. Development 141, 3458-71 (2014).

3. Luteijn, M.J. & Ketting, R.F. PIWI-interacting RNAs: from generation to transgenerational epigenetics. Nat Rev Genet 14, 523-34 (2013).

4. Czech, B. & Hannon, G.J. One Loop to Rule Them All: The Ping-Pong Cycle and piRNA-Guided Silencing. Trends Biochem Sci 41, 324-337 (2016).

5. Luteijn, M.J. et al. Extremely stable Piwi-induced gene silencing in Caenorhabditis elegans. EMBO J 31, 3422-30 (2012).

6. Sakakibara, K. & Siomi, M.C. The PIWI-Interacting RNA Molecular Pathway: Insights From Cultured Silkworm Germline Cells. Bioessays 40(2018).

7. Rojas-Rios, P. & Simonelig, M. piRNAs and PIWI proteins: regulators of gene expression in development and stem cells. Development 145(2018).

8. Das, P.P. et al. Piwi and piRNAs act upstream of an endogenous siRNA pathway to suppress Tc3 transposon mobility in the Caenorhabditis elegans germline. Mol Cell 31, 79-90 (2008).

9. Batista, P.J. et al. PRG-1 and 21U-RNAs interact to form the piRNA complex required for fertility in C. elegans. Mol Cell 31, 67-78 (2008).

10. Wang, G. & Reinke, V. A C. elegans Piwi, PRG-1, regulates 21U-RNAs during spermatogenesis. Curr Biol 18, 861-7 (2008).

11. Bagijn, M.P. et al. Function, targets, and evolution of Caenorhabditis elegans piRNAs. Science 337, 574-578 (2012).

12. Lee, H.C. et al. C. elegans piRNAs mediate the genome-wide surveillance of germline transcripts. Cell 150, 78-87 (2012).

13. Simon, M. et al. Reduced insulin/IGF-1 signaling restores germ cell immortality to Caenorhabditis elegans Piwi mutants. Cell Rep 7, 762-73 (2014).

14. Heestand, B., Simon, M., Frenk, S., Titov, D. & Ahmed, S. Transgenerational Sterility of Piwi Mutants Represents a Dynamic Form of Adult Reproductive Diapause. Cell Rep 23, 156-171 (2018).

15. de Albuquerque, B.F., Placentino, M. & Ketting, R.F. Maternal piRNAs Are Essential for Germline Development following De Novo Establishment of Endo-siRNAs in Caenorhabditis elegans. Dev Cell 34, 448-56 (2015).

16. Phillips, C.M., Brown, K.C., Montgomery, B.E., Ruvkun, G. & Montgomery, T.A. piRNAs and piRNA-Dependent siRNAs Protect Conserved and Essential C. elegans Genes from Misrouting into the RNAi Pathway. Dev Cell 34, 457-65 (2015).

17. Ashe, A. et al. piRNAs can trigger a multigenerational epigenetic memory in the germline of C. elegans. Cell 150, 88-99 (2012).

18. Shirayama, M. et al. piRNAs initiate an epigenetic memory of nonself RNA in the C. elegans germline. Cell 150, 65-77 (2012).

19. Buckley, B.A. et al. A nuclear Argonaute promotes multigenerational epigenetic inheritance and germline immortality. Nature 489, 447-51 (2012).

20. Ruby, J.G. et al. Large-scale sequencing reveals 21U-RNAs and additional microRNAs and endogenous siRNAs in C. elegans. Cell 127, 1193-207 (2006).

21. Kasper, D.M., Wang, G., Gardner, K.E., Johnstone, T.G. & Reinke, V. The C. elegans SNAPc component SNPC-4 coats piRNA domains and is globally required for piRNA abundance. Dev Cell 31, 145-58 (2014).

22. Weick, E.M. et al. PRDE-1 is a nuclear factor essential for the biogenesis of Ruby motif-dependent piRNAs in C. elegans. Genes Dev 28, 783-96 (2014).

23. Beltran, T. et al. Evolutionary analysis im[licates RNA polymersase II pausing and chromatin structure in nematode piRNA biogenesis. BioRxiv https://doi.org/10.1101/281360 (2018).

24. Blumenthal, T. Trans-splicing and operons in C. elegans. WormBook, 1-11 (2012).

25. Gu, W. et al. CapSeq and CIP-TAP identify Pol II start sites and reveal capped small RNAs as C. elegans piRNA precursors. Cell 151, 1488-500 (2012).

26. de Albuquerque, B.F. et al. PID-1 is a novel factor that operates during 21U-RNA biogenesis in Caenorhabditis elegans. Genes Dev 28, 683-8 (2014).

27. Goh, W.S. et al. A genome-wide RNAi screen identifies factors required for distinct stages of C. elegans piRNA biogenesis. Genes Dev 28, 797-807 (2014).

28. Tang, W., Tu, S., Lee, H.C., Weng, Z. & Mello, C.C. The RNase PARN-1 Trims piRNA 3’ Ends to Promote Transcriptome Surveillance in C. elegans. Cell 164, 974-84 (2016).

29. Kamminga, L.M. et al. Differential impact of the HEN1 homolog HENN-1 on 21U and 26G RNAs in the germline of Caenorhabditis elegans. PLoS Genet 8, e1002702 (2012).

30. Billi, A.C. et al. The Caenorhabditis elegans HEN1 ortholog, HENN-1, methylates and stabilizes select subclasses of germline small RNAs. PLoS Genet 8, e1002617 (2012).

31. Montgomery, T.A. et al. PIWI associated siRNAs and piRNAs specifically require the Caenorhabditis elegans HEN1 ortholog henn-1. PLoS Genet 8, e1002616 (2012).

32. Cecere, G., Zheng, G.X., Mansisidor, A.R., Klymko, K.E. & Grishok, A. Promoters recognized by forkhead proteins exist for individual 21U-RNAs. Mol Cell 47, 734-45 (2012).

33. Sugiyama, T. et al. Enhancer of Rudimentary Cooperates with Conserved RNA-Processing Factors to Promote Meiotic mRNA Decay and Facultative Heterochromatin Assembly. Mol Cell 61, 747-759 (2016).

34. Jankowska-Anyszka, M. et al. Multiple isoforms of eukaryotic protein synthesis initiation factor 4E in Caenorhabditis elegans can distinguish between mono-and trimethylated mRNA cap structures. J Biol Chem 273, 10538-42 (1998).

35. Arai, R. et al. Crystal structure of an enhancer of rudimentary homolog (ERH) at 2.1 Angstroms resolution. Protein Sci 14, 1888-93 (2005).

36. Frokjaer-Jensen, C. et al. Random and targeted transgene insertion in Caenorhabditis elegans using a modified Mos1 transposon. Nat Methods 11, 529-34 (2014).

37. Sengupta, M.S. et al. ifet-1 is a broad-scale translational repressor required for normal P granule formation in C. elegans. J Cell Sci 126, 850-9 (2013).

38. Cauchi, R.J. SMN and Gemins: ‘we are family’ … or are we?: insights into the partnership between Gemins and the spinal muscular atrophy disease protein SMN. Bioessays 32, 1077-89 (2010).

39. Mader, S., Lee, H., Pause, A. & Sonenberg, N. The translation initiation factor eIF-4E binds to a common motif shared by the translation factor eIF-4 gamma and the translational repressors 4E-binding proteins. Mol Cell Biol 15, 4990-7 (1995).

40. Gruner, S. et al. The Structures of eIF4E-eIF4G Complexes Reveal an Extended Interface to Regulate Translation Initiation. Mol Cell 64, 467-479 (2016).

41. Minasaki, R., Puoti, A. & Streit, A. The DEAD-box protein MEL-46 is required in the germ line of the nematode Caenorhabditis elegans. BMC Dev Biol 9, 35 (2009).

42. Keiper, B.D. et al. Functional characterization of five eIF4E isoforms in Caenorhabditis elegans. J Biol Chem 275, 10590-6 (2000).

43. Maris, C., Dominguez, C. & Allain, F.H. The RNA recognition motif, a plastic RNA-binding platform to regulate post-transcriptional gene expression. FEBS J 272, 2118-31 (2005).

44. Schirle, N.T. & MacRae, I.J. The crystal structure of human Argonaute2. Science 336, 1037-40 (2012).

45. Siomi, M.C., Mannen, T. & Siomi, H. How does the royal family of Tudor rule the PIWI-interacting RNA pathway? Genes Dev 24, 636-46 (2010).

46. Philippe, L. et al. An in vivo genetic screen for genes involved in spliced leader trans-splicing indicates a crucial role for continuous de novo spliced leader RNP assembly. Nucleic Acids Res 45, 8474-8483 (2017).

47. Xu, C. et al. Structural insights into Gemin5-guided selection of pre-snRNAs for snRNP assembly. Genes Dev 30, 2376-2390 (2016).

48. Zanetti, S., Meola, M., Bochud, A. & Puoti, A. Role of the C. elegans U2 snRNP protein MOG-2 in sex determination, meiosis, and splice site selection. Dev Biol 354, 232-41 (2011).

49. Ferguson, K.C., Heid, P.J. & Rothman, J.H. The SL1 trans-spliced leader RNA performs an essential embryonic function in Caenorhabditis elegans that can also be supplied by SL2 RNA. Genes Dev 10, 1543-56 (1996).

50. Edgar, R.C. MUSCLE: multiple sequence alignment with high accuracy and high throughput. Nucleic Acids Res 32, 1792-7 (2004).

51. Robert, X. & Gouet, P. Deciphering key features in protein structures with the new ENDscript server. Nucleic Acids Res 42, W320-4 (2014).

